# Not such silly sausages: Northern quolls exhibit aversion to toads after training with toad sausages

**DOI:** 10.1101/181149

**Authors:** Naomi Indigo, James Smith, Jonathan K. Webb, Ben Phillips

**Affiliations:** School of Life Sciences, University of Technology Sydney, PO Box 123 Broadway NSW 2007. Australia. E; Australian Wildlife Conservancy, Mornington Wildlife Sanctuary. PMB 925, Derby, WA 6728, Australia. E; School of Biosciences, University of Melbourne. School of BioSciences, Royal Parade, Parkville, Victoria 3010, Australia. E

**Author notes:** Corresponding Author: Naomi Indigo, E.

**Keywords:** conditioned taste aversion, invasive species, predator–prey, *Rhinella marina*, *Dasyurus hallucatus*, bait uptake, target specificity, behavioural change, *Bufo marinus*

## Abstract

The invasion of toxic cane toads (*Rhinella marina*) is a major threat to northern quolls (*Dasyurus hallucatus*) which are poisoned when they attack this novel prey item. Quolls are now endangered as a consequence of the toad invasion. Conditioned taste aversion can be used to train individual quolls to avoid toads, but we currently lack a training technique that can be used at a landscape scale to buffer entire populations from toad impact. Broad scale deployment requires a bait that can be used for training, but there is no guarantee that such a bait will ultimately elicit aversion to toads. Here we test a manufactured bait—a ‘toad sausage’—for its ability to elicit aversion to toads in northern quolls. To do this, we exposed one group of quolls to a toad sausage and another to a control sausage and compared the quolls’ predatory responses when presented with a dead adult toad. Captive quolls that consumed a single toad sausage showed substantially reduced interest in cane toads, interacting with them for less than half the time of their untrained counterparts and showing substantially reduced attack behaviour. We also quantified bait uptake in the field, by both quolls and non-target species. These field trials showed that wild quolls were the most frequent species attracted to the baits, and that approximately 61% of quolls consumed toad-aversion baits when first encountered. Between 40-68% of these animals developed aversion to further bait consumption. Our results suggest that toad-aversion sausages can be used to train wild quolls to avoid cane toads. This opens the possibility for broad-scale quoll training with toad aversion sausages: a technique that may allow wildlife managers to prevent quoll extinctions at a landscape scale.

## Introduction

Invasive species are a major threat to biodiversity (Reaser *et al.* 2007; Woinarski, Burbidge & Harrison 2014). In Australia, species such as feral cats (*Felis catus*) (Legge *et al.* 2017), domestic dogs (*Canis familiaris*) (Doherty *et al.* 2017), foxes (*Vulpes vulpes*) (Short & Smith 1994; Risbey *et al.* 2000) and cane toads (*Rhinella marina*) (Burnett 1997; Letnic, Webb & Shine 2008; Jolly, Shine & Greenlees 2015) all have serious impacts on native species. Controlling these species at a landscape scale, however, has proved extremely difficult (Ziembicki *et al.* 2015). Because of this, increasing attention is being paid to mitigating the impact of invasives, rather than supressing their populations (Simberloff *et al.* 2013).

Cane toads are a case in point. These invasive amphibians now occupy more than 1.5 million square kilometres of Australia, continue to spread (Urban *et al.* 2007), and have proved difficult to control. The cane toads’ defensive chemicals (bufadienalides and related toxins) are highly cardioactive and are unlike toxins possessed by native Australian animals (Hayes *et al.* 2009). As a result, many vertebrate predators, including varanid lizards, snakes, and marsupial predators such as quolls, die after attacking or consuming toads (Covacevich & Archer 1975; Webb, Shine & Christian 2005; Smith & Phillips 2006; Hayes *et al.* 2009; Shine 2010). Some reptilian predator populations have adapted to the presence of toads by evolving innate aversion to toads (Phillips & Shine 2005; Llewelyn *et al.* 2011). In the short term, some marsupial predators rapidly learn to avoid toads as prey (Webb *et al.* 2008; Webb, Pearson & Shine 2011). An obvious avenue for mitigating the impact of toads, then, is to train predators to avoid toads (Webb *et al.* 2008).

Such training can be achieved through conditioned taste aversion (CTA). Conditioned taste aversion is a powerful innate response found across all vertebrates; an evolved defence mechanism against poisoning (Sinclair & Bird 1984; Conover 1995; Cohn & MacPhail 1996; Bernstein 1999; Mappes, Marples & Endler 2005; Page & Ryan 2005; Glendinning 2007). With CTA, animals acquire an aversion to a referent food as a result of a nauseating experience (Gustavson & Nicolaus 1987). Agriculturalists and wildlife managers have used conditioned taste aversion to reduce wildlife damage to crops, industry, or livestock (Gustavson *et al.* 1974; Ellins & Catalano 1980; Avery 1985; Provenza *et al.* 1990; Ternent & Garshelis 1999; Smith *et al.* 2000). CTA has also been used successfully to reduce predation on native or introduced wildlife (Nicolaus & Nellis 1987; Conover 1989; Nicolaus *et al.* 1989; Semel & Nicolaus 1992; Avery *et al.* 1995; Bogliani & Fiorella 1998; Cox *et al.* 2004).

One of the Australian species most strongly impacted by cane toads is the northern quoll, *Dasyurus hallucatus*. As toads have spread, they have caused numerous local extinctions of this native marsupial predator (Burnett 1997; Oakwood & Foster 2008). CTA training using small toads infused with the nausea inducing chemical thiabendazole (TBZ) elicits aversion to live toads in northern quolls (O'Donnell, Webb & Shine 2010), suggesting the technique has promise as a management tool for mitigating toad impact. Capacity to elicit aversion is, however, only the first hurdle. To be effective as a management tool, CTA needs to meet two additional conditions. First, CTA training needs to be deliverable to a large number of individuals under field conditions. Second, prey aversion needs to occur in a large enough proportion of the population, and be behaviourally persistent for long enough (within and across generations), that population-level benefits are realised. In quolls, it is clear that CTA training in captivity can be used to elicit toad aversion, and that this aversion improves survival rates when animals are released into the field (O'Donnell, Webb & Shine 2010). More importantly, parentage analyses demonstrated that some offspring of surviving ‘toad smart’ females also survived and reproduced (Cremona *et al.* 2017), suggesting that training a single generation could yield significant conservation benefits. The remaining challenge then is to effectively deliver CTA training to a large number of individuals under field conditions.

In captivity, CTA training was achieved by feeding quolls a small non-lethal-sized toad laced with the nausea-inducing chemical thiabendazole. Such a strategy is not feasible at a large scale in a field setting. To achieve in situ training at scale requires use of a manufactured training bait. Any bait, of course, needs to fulfil the criteria we have identified above: elicits aversion to toads, has a high uptake rate; and effectively trains a high enough proportion of the population that population persistence is assured. An additional consideration is whether the bait is taken by non-target species. This is a major concern in lethal baiting campaigns (Sinclair & Bird 1984; Avery *et al.* 1995; Fairbridge *et al.* 2003; Glen & Dickman 2003; Claridge & Mills 2007; Jolley *et al.* 2012), but a smaller consideration in non-lethal baiting such as we envisage here. Non-target uptake remains important, however, because it can reduce target species’ access to bait and so significantly increase the cost and complexity of the baiting effort. Because of this, it is important to understand non-target species uptake rates.

In this study we assess the value of a manufactured bait (‘toad aversion sausages’). We ask whether quolls generalise their CTA from the bait to toads, whether the bait is taken up by wild quolls (and non-target species), and whether it appears to elicit CTA under field conditions.

## Methods

### Cane toad sausages

Cane toad sausages were made up of 15g of minced skinned adult cane toad legs, 1 whole cane toad metamorph weighing <2g, and 0.06g of Thiabendazole (per sausage; dose rate less than 300mg/kg adult quoll body weight, determined by the smallest – 200g – adult seen at our study site) packed into a synthetic sausage skin. In our captive trials, we used the same sausage composition, to accurately reflect our field scenario. Thiabendazole is an inexpensive, broad-spectrum anthelmintic and antifungal agent (Robinson, Stoerk & Graessle 1965). It is orally-effective and regarded as relatively safe, producing low mammalian mortality: oral LD_50_ is 2.7g/kg body weight (Dilov *et al.* 1981). It is fast acting and peak concentration occurs in the plasma one hour after consumption (Tocco *et al.* 1966). Thiabendazole has produced strong aversions to treated foods in lab rats (Gill, Whiterow & Cowan 2000; Massei & Cowan 2002), wolves (*Canis lupus*) (Gustavson, Gustavson & Holzer 1983; Ziegler *et al.* 1983), and black bears (*Ursus americanus*) (Ternent & Garshelis 1999). Thiabendazole induces a robust CTA after a single oral dose (Nachman & Ashe 1973; O'Donnell, Webb & Shine 2010) and is physically stable at ambient conditions in the bait substrate (Gill, Whiterow & Cowan 2000; Massei, Lyon & Cowan 2003).

### Captive trials

The uptake of toad aversion sausages by *D.hallucatus* and their subsequent response to toads was observed in captive northern quolls previously collected from toad-free areas of Astell Island, and then housed at the Territory Wildlife Park, Northern Territory. Animals (9 male and 9 female) were randomly allocated treatment (*n=* 9) or control (*n=*9) sausage groups. Treatment sausages were exactly as described previously. Control sausages were comprised of store purchased beef sausages. These were selected as a control sausage as it was an item that animals are also not familiar with to control for hunger differences and any possible neophobic responses.

To measure individual responses to cane toads following ingestion of sausage, each individual was presented with a dead adult cane toad the following evening. The dead adult toad was secured in a wire cage, so that animals could see and smell the prey item but not access it. The experiment was run over 3 nights. Experiments began at sunset and ran for on average 2 hours. The response was filmed using a GoPro Hero 3 White camera (GoPro Inc, San Mateo, California, USA).

### Field trials

#### Study area

The field study was conducted between May 2016-February 2017 at Mornington Wildlife Sanctuary, a 300,000 ha property in the central Kimberley region of western Australia managed for conservation by the Australian Wildlife Conservancy (17°01’S, 126°01’E; Fig. 1). The area is characterized by savanna woodland dissected by sandstone gorges of varying topographic complexity. On average, this area receives 788 mm of rain annually, most of which falls during the wet season from November to April.

**Figure 1:**
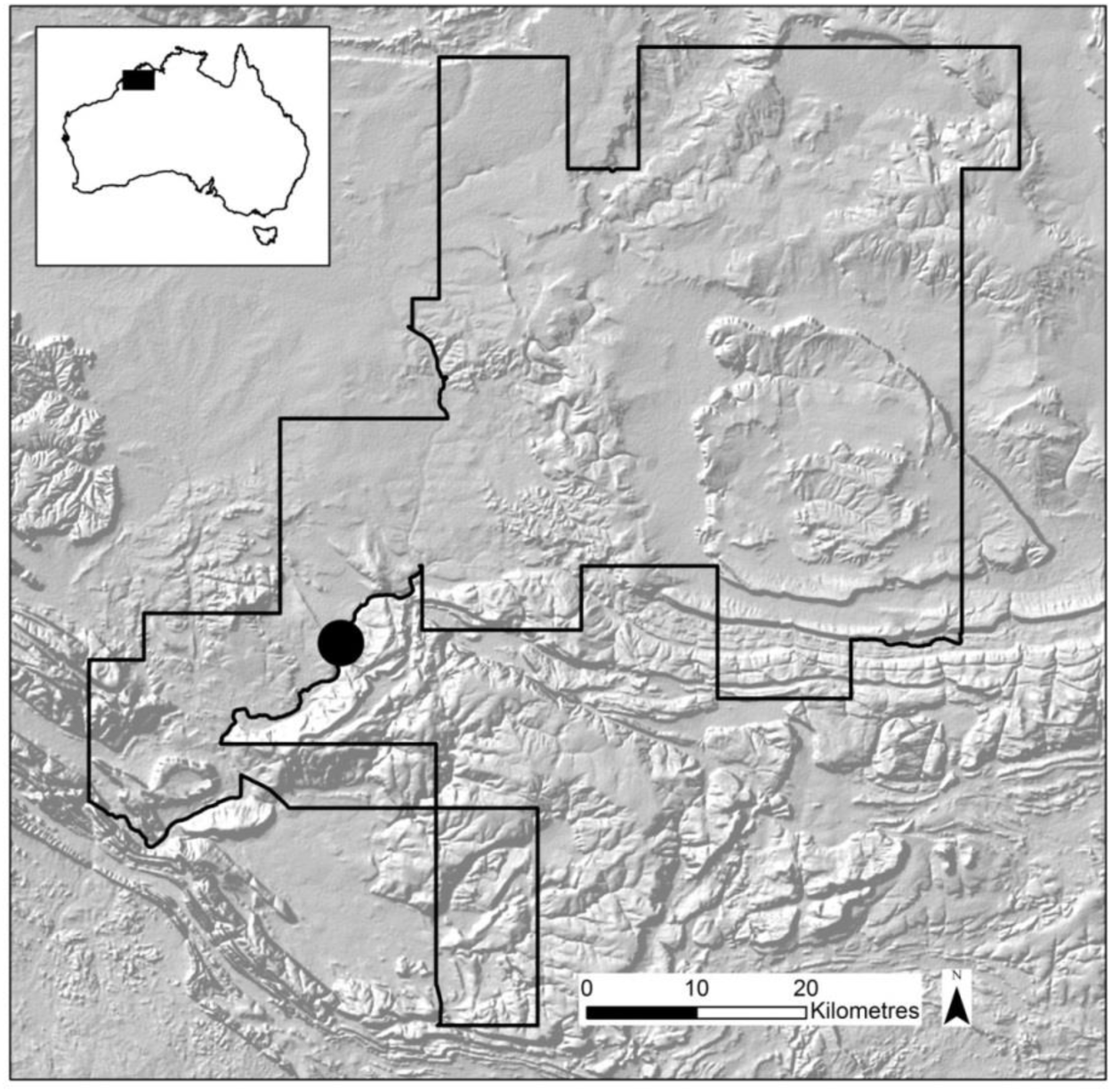
Location of the study area within Australian Wildlife Conservancy’s Mornington Wildlife Sanctuary, in the central Kimberley, Western Australia.

We worked at four sites on the property; Site 1 (SJ) was at Sir John Gorge (17°31.780S, 126°13.080E) along the Fitzroy River. Site 2 (KP) (17°31’43.032, 126°13’11.050) was approximately 2 km upstream from Site 1 in the same gorge. Site 3 (TC) (17° 30’ 37.213,126°14’4.092) was 5 km upstream from Site 2 in a narrow rocky gorge that feeds into Sir John Gorge. Site 4 (RP) (17°35’12.119, 126°19’21.959) was a narrowly incised sandstone gorge following a watercourse within rocky range country approximately 9 km north-east of Site 1. Sites were selected based on the detection of quolls in the Australian Wildlife Conservancy’s fauna surveys (AWC unpub. data). At the time of the study, toads were yet to arrive at our sites; they subsequently arrived by March 2017.

#### CTA sausage field trials

In this study, “site” is the location where an experiment took place. “Bait station” is a location within a site where sausage bait was offered. A “session”, is a time interval when bait stations were active. A total of four sessions were conducted approximately five months apart. Sessions recorded up to four “bait events”. Bait Events are defined as an occasion when new bait was placed at a bait station and (if still existing) the old bait removed.

Each site contained 20 bait stations placed 50-80m apart in a linear transect along a gorge wall where the presence of *D. hallucatus* was previously confirmed (AWC, unpub. data). Bait stations consisted of one cane toad sausage placed under a single camera trap (White flash and Infrared Reconyx Motion Activated, (HP800, U.S.A). Cameras were secured to trees or rocky ledges approximately 1m from the ground and aligned to face directly downwards (Diete *et al.* 2016). Cameras were set to take five consecutive photographs for each trigger with no delay between triggers. Each cane toad sausage was placed inside a ring of powdered insecticide (Coopex) to protect from ant spoilage. Each session’s bait stations were rebaited up to three times (for a maximum of 4 bait events within any given session) whereby bait stations were rebaited with a fresh CTA bait and the old bait removed (Table 1). A total of 513 individual cane toad sausages were deployed over the period of study.

**Table 1:** CTA sessions and bait events, * denotes empty cells. Bait events occurred at the same time within each site. KP was baited only once in November to expand the sample size and CTA train quolls prior to cane toad arrival.

| Site Name | Session Year | Session Month | No. Bait Events (BE) | BE 1-date | BE 2-date | BE 3-date | BE 4-date | No. of bait stations<sup>600</sup> |
| --- | --- | --- | --- | --- | --- | --- | --- | --- |
| KP | 2016 | Nov | 1 | 31/10/16 | 1/11/16 | 2/11/16 | * | 20 |
| RP | 2017 | Feb | 1 | 3/2/2017 | * | * | * | 20 |
| RP | 2016 | May | 3 | 10/5/16 | 13/5/16 | 21/5/16 | * | 20 |
| RP | 2016 | Sept | 3 | 15/9/16 | 16/9/16 | 17/9/16 | * | 20 |
| SJ | 2017 | Feb | 1 | 3/2/2017 | * | * | * | 20 |
| SJ | 2016 | May | 3 | 10/5/16 | 13/5/16 | 21/5/16 | * | 20 |
| SJ | 2016 | Sept | 4 | 15/9/16 | 16/9/16 | 17/9/16 | 19/9/16 | 20 |
| TC | 2017 | Feb | 1 | 3/2/2017 | * | * | * | 33 |
| TC | 2016 | May | 3 | 10/5/16 | 13/5/16 | 21/5/16 | * | 20 |
| TC | 2016 | Sept | 3 | 15/9/16 | 16/9/16 | 17/9/16 | * | 20 |

### Data analysis

#### Captive trials

Videos were scored by the same observer who was blind to the quoll’s treatment or control group. Following Kelly and Phillips (2017) we separated the time that quolls spent exhibiting various predatory behaviours into three categories: “Sniff”, “Investigate” and “Attack”. Sniff was defined as when quolls were visibly twitching their nose in the direction of the toad, “Investigating” behaviour was defined as the quoll being engaged with the cage containing the toad, exhibiting scent marking or digging around the outside of cane toad enclosure and “attack” behaviour was defined as quolls exhibited pawing or licking or biting behaviour to toads cages. We summed all of these to measure the total time spent interacting with a toad. We converted each of these variables to a proportion of time spent in each of these activities, where the denominator was the total time that the animal was observable on camera. These response variables were not normally distributed, and could not be made to conform to normality through transformation. Because of this we used bootstrapping to obtain confidence intervals for the mean time engaged in each behaviour, and to test the null hypothesis that there was no difference between treatments in mean time spent in each activity. The perception that animals exhibit a lower propensity towards attacking a prey item following ingestion and subsequent malaise during CTA training is non-controversial (Gustavson *et al.* 1974; Gustavson *et al.* 1976; Gustavson 1982; Gustavson & Basche 1983; Gustavson, Gustavson & Holzer 1983; Ziegler *et al.* 1983; Gustavson & Nicolaus 1987; Nicolaus 1987; Nicolaus & Nellis 1987; Nicolaus *et al.* 1989; Schneider & Pinnow 1994; Smith *et al.* 2000; Riley & Freeman 2004; Sevelinges *et al.* 2009; O'Donnell, Webb & Shine 2010; Thornton & Raihani 2010; Thornton & Clutton-Brock 2011). More relevant to this study is the outcomes of previous trails by O'Donnell, Webb and Shine (2010) and Kelly and Phillips (2017) where quolls exhibited less interest in prey items after consuming a toad metamorph laced with thiabendazole. Based on these previous results, we had a strong *a priori* expectation that animals could either be unaffected or only become less interested in toads after ingestion of cane toad sausages. Thus, we employed a one-tailed test, with the alternative hypothesis that the mean time spent investigating and attacking toads will be lower in the treatment group. This analysis was performed using R (R Core Team, 2016).

#### Field trials

Images from bait stations were collated and tagged by pass, session, site, bait-event, species and activity. A ‘pass’ was defined as when a new species entered the frame or when images that were at least 5 minutes between when the previous detection of the same species passed. This reduced any likelihood of individuals of the same species being overlooked during analysis. “Activity” was hierarchical, with the highest activity being ‘Bait taken’; this was defined as either photographic evidence of animal eating bait or bait being taken from the bait station. ‘Bait investigated’ was defined as when bait was sniffed but not consumed or taken. ‘Bait area investigated with no bait available’ was defined as when no bait was available at a bait station, but the animal was still visiting or investigating the bait station.

We analysed data using two levels of observation to determine 1) which species were attracted to bait, and 2) which species took bait. A frequency distribution (*n* times each species was recorded) was calculated and the proportion of bait takers in each species was estimated. Passes in which we were unable to identify the species were pooled and removed from further analysis. Additionally, if a species total number of visits was less than 10 we removed that species from the analysis. Additionally *Varanus tristis, V. panoptes, V. mitchelli* and *V. mertensi* were pooled into ‘*Varanus* other species’ due to small sample sizes. We identified individual *D. hallucatus* that visited bait stations by their unique spot patterns (Hohnen *et al.* 2013) to determine visitation rate and bait uptake of individuals. To do this we employed Wild ID (Version 1.0, January 2011) (Bolger *et al.* 2011) to extract distinctive image features in animals spot patterns, the program calculates a matching score that characterizes the goodness of fit between two images. These matching scores were then used to rank and select matches to each focal image. We also conducted manual checks with all photographs and compared them to those already identified to determine whether a new individual had been recorded. Quolls were identified to individual within each session, and we treat each session (separated by a minimum of four months) as independent with regard to quoll ID and behaviour. This decision was made for logistical reasons (difficulty of identifying individuals using spot ID), but is supported by exploratory analysis of first pass uptake rates showing that these do not vary systematically with session (see Results). It is likely, therefore, that any training is forgotten within the 4-5 month window between sessions.

## Results

### Captive trials

Of the treatment animals, seven (77%) consumed all or part of a cane toad sausage and eight (88%) control animals consumed beef sausages. Treatment had no significant effect on whether the initial sausage was consumed, (χ^2^= 0.0, df = 1, *p* = 1). In our video trials, quolls spent an average of only 0.6% of the total time on camera interacting with the toad. Control animals, however, spent more than twice as much time interacting with the toad than treatment animals (control = 0.95%; treatment = 0.42%, bootstrap p-value = 0.022). When we break this down by specific types of interaction, control animals spend approximately sixty times longer investigating (control = 0.15%; treatment = 0.00024%, bootstrap p-value = 0.051); twice as much time sniffing (control = 0.70%; treatment = 0.35%, bootstrap p-value = 0.044); and twenty times more time attacking (control = 0.03%; treatment = 0.0015%, bootstrap p-value = 0.036) toads when compared with the control (Figure 2).

**Figure 2:**
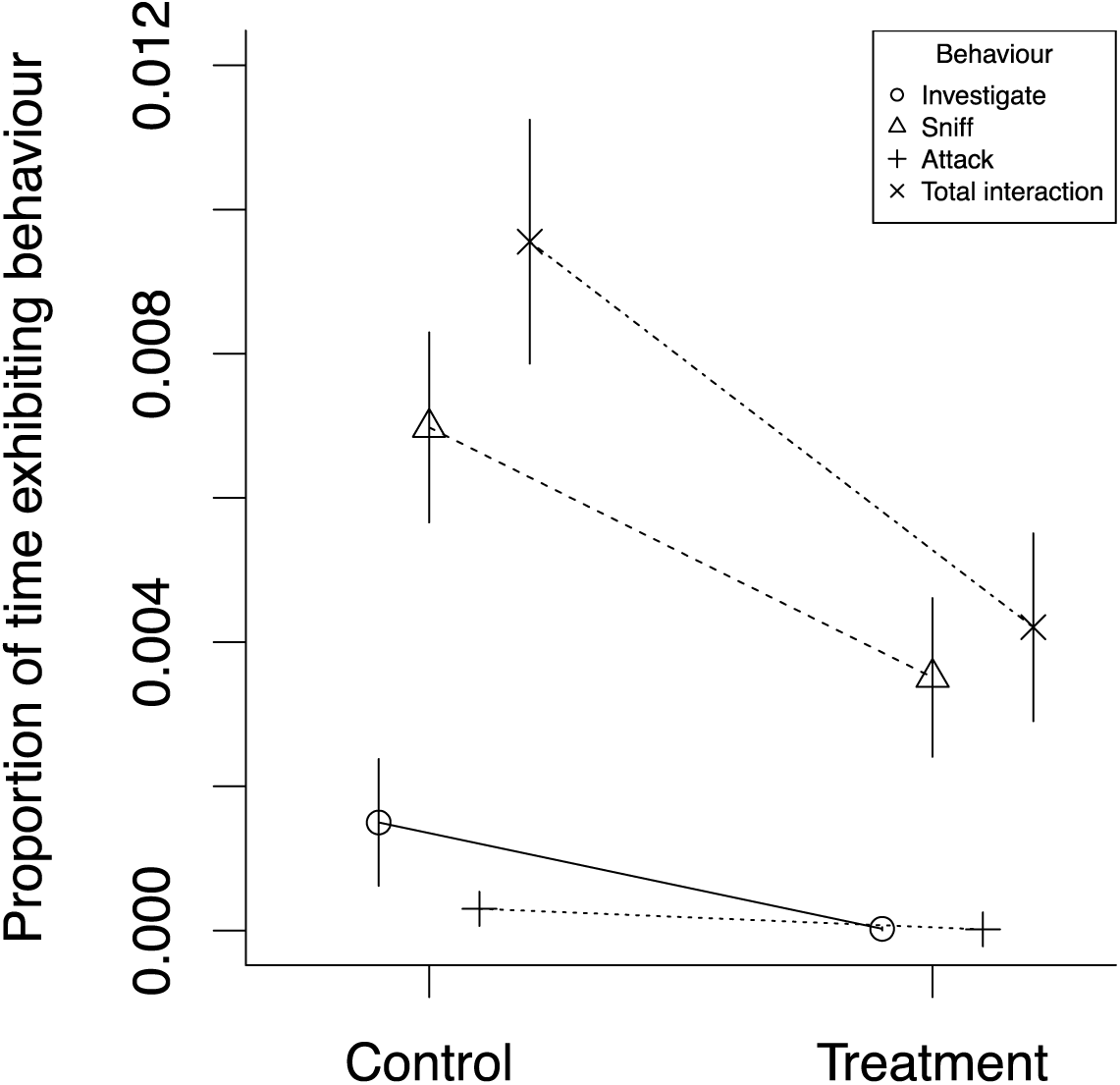
Mean proportion of active time that quolls spent directed towards toads. Behaviours are split into categories and across control and treatment groups. Error bars represent bootstrap standard errors.

### Field trials

#### Target and non-target uptake

A total of 26 species were captured on camera traps visiting bait stations. For eleven of these species, there were sufficient data to compare their response to bait uptake. The most frequent visitors to the bait stations were quolls, with *n =* 345 passes (Figure 3). Almost all bait removal was executed by quolls that took 65 baits of the 90 baits removed. Other species took far fewer: *Zyzomys argurus,* 9; *Ctenotus spp*., 2; *Pseudantechinus ningbing*, 2; *Varanus glauerti*, 2; and *Varanus glebopalma*, 2.

**Figure 3:**
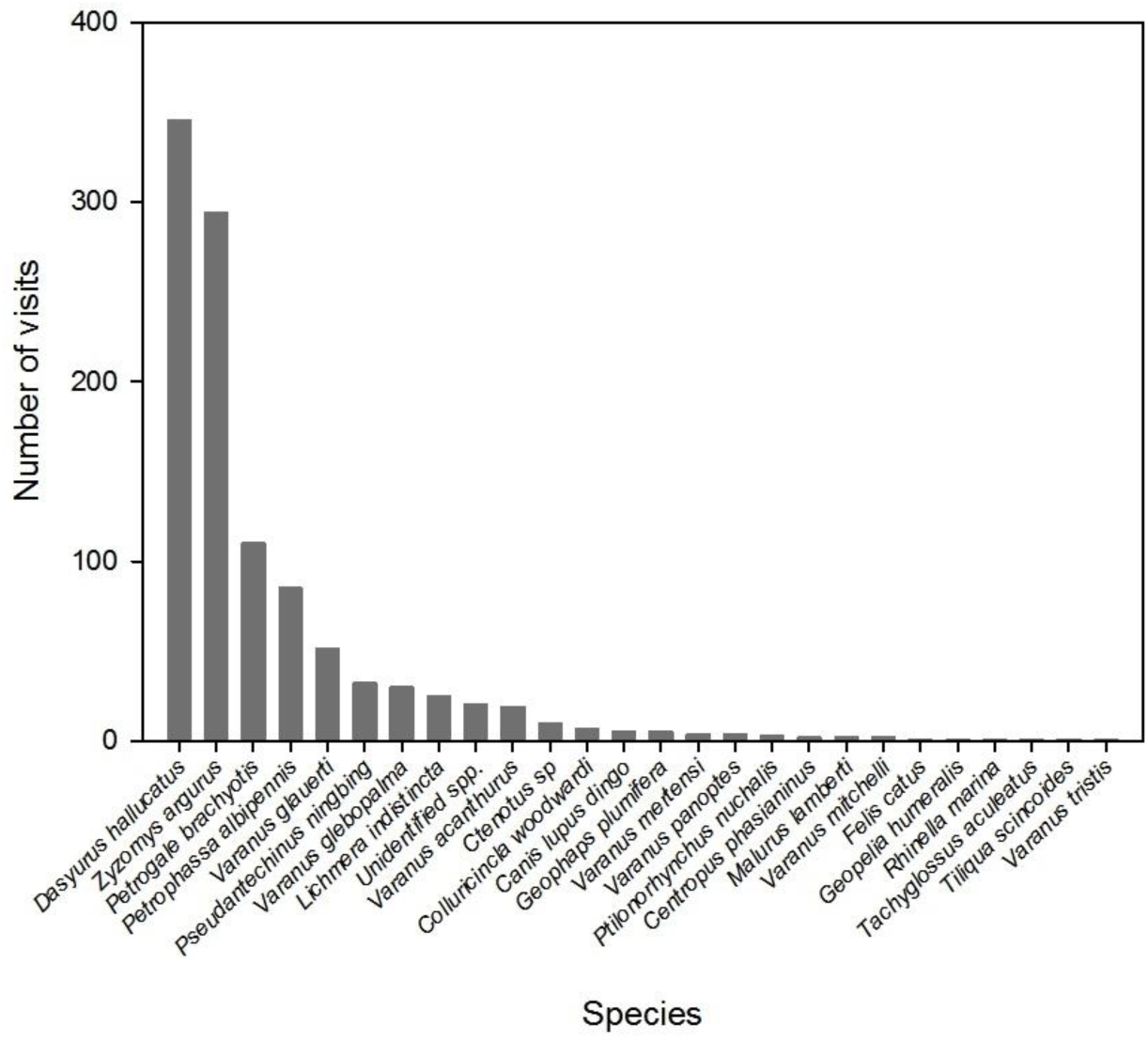
Frequency of visits to CTA bait stations by each species. Unidentified species group comprises unidentified rodents, birds, and frogs.

#### Target uptake and training rates

First pass uptake responses to the bait did not vary systematically across sessions (χ^2^=1.7, df=4, p=0.79; Fig. 4). We thus treated individuals as independent across sessions with regard to behaviour.

**Figure 4:**
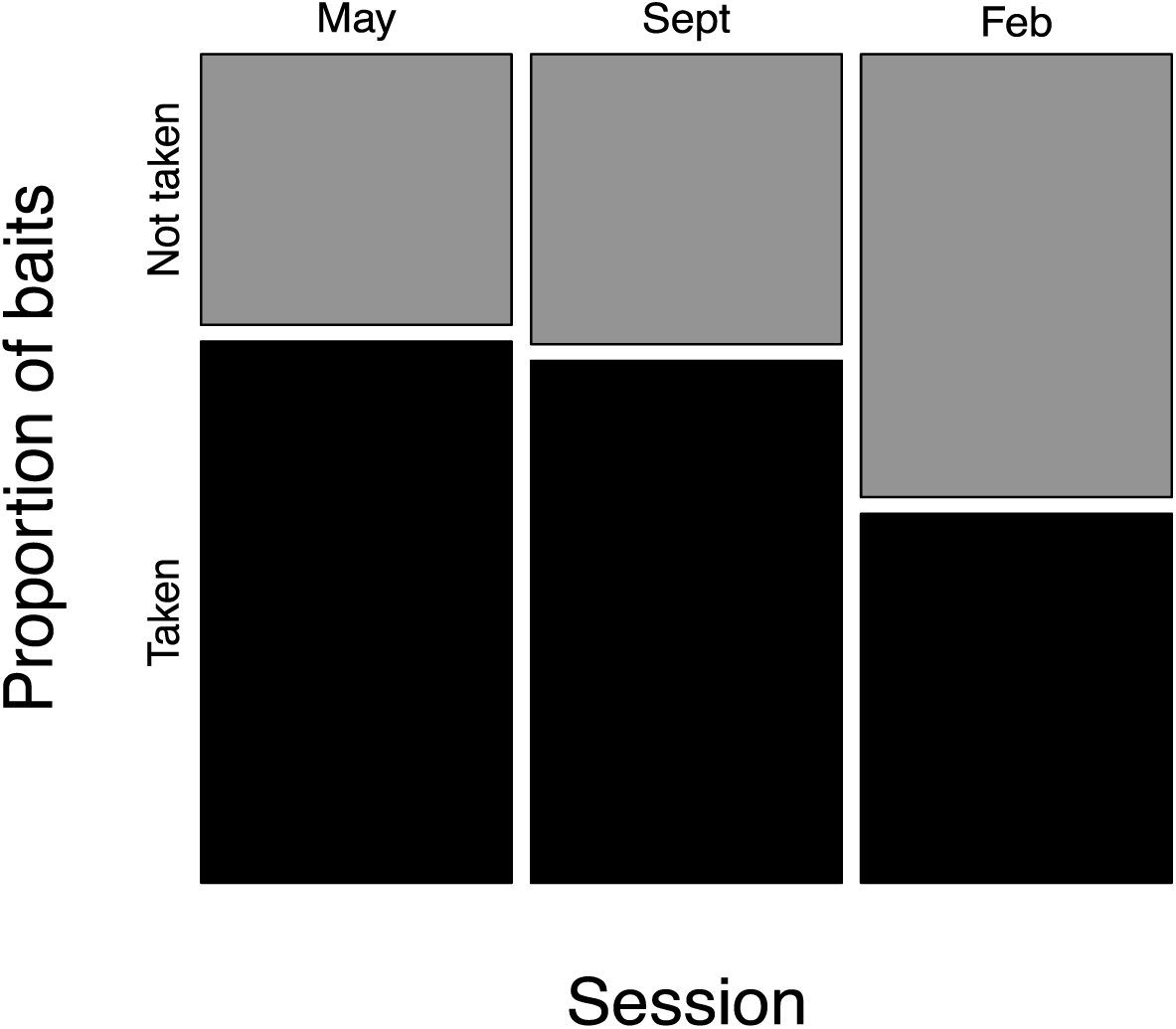
First pass behavioural responses of northern quolls to the bait each session. “Sept” includes the one November session also.

Following identification of individual quolls within sessions, it became apparent that bait stations were visited by a total of 70 “individual” quolls over the period of the study. Of these individuals, and considering only their first bait encounter, the bait was taken by 28 individuals and rejected (bait investigated but not taken) by 18 individuals. Thus the total bait uptake rate at first encounter was 61% (SE=7.2%). A further 24 individuals only visited bait stations when there was no bait available.

Across all passes, a total of 31 quolls consumed a bait. Ten of these animals ultimately consumed baits on more than one occasion within a session (32%, SE=8.3%). Clearly, these individuals were not effectively trained, failing to even exhibit aversion to the bait. We have two ways of estimating the conversion rate (from untrained to trained, given bait consumption). Placing an upper bound, we could consider all individuals that took a bait but were not observed to take a second bait (20 of 31 = 68%) as trained. For a lower bound, we could take the conservative approach and consider only those known to have consumed a bait and then seen to approach and reject a bait as trained (7 of 17 = 41%). Thus, somewhere between 41 and 68% of animals consuming a bait appear to have been trained.

## Discussion

Our captive trials clearly indicate that training a quoll using a thiabendazole-laced toad sausage changes their behaviour towards toads. Although our sample sizes were modest and not all of our treatment animals fully consumed the bait, it is clear that sausage-trained animals spent substantially less time interacting with a toad—between one half to one sixtieth of the time as control animals. This behavioural shift is reflected across all prey acquisition behaviours: investigating, sniffing, and attacking. Quolls clearly generalise their acquired aversion from the bait to a real toad.

The field trials show that the toad sausages are attractive to quolls. Although 26 species encountered the baits, quolls were the most frequent visitors to the bait at our study sites, and were far and away the most likely species to consume the bait. Thus, non-target uptake is relatively modest, compared with the high level of uptake of baits by non-target animals observed in other lethal-baiting studies (Cowled *et al.* 2006; Dundas, Adams & Fleming 2014) It is more difficult to estimate the rate of successful training in the field, but it is likely that between 41-68% of animals consuming a bait in the field have been successfully trained. The apparent independence of quoll behaviour to bait uptake across sessions also suggests that, in the absence of further reinforcing stimulus (i.e., cane toads), CTA training potentially only elicits aversion for a limited time (<4 months).

The TBZ dose of <300mg/kg animal body weight in our cane toad sausages was relatively low compared to earlier work (400mg/kg in (Jolly et al.unpublished data; O'Donnell, Webb & Shine 2010; Cremona *et al.* 2017; Kelly & Phillips 2017) but was set low by regulators (Australian Pesticides and Veterinary Medicines Authority-(APVMA)) to allow for potential multiple bait uptake, sub-adult target, and non-target species. Given the LD_50_ of TBZ is more than nine times higher than our dose rate; the delivered dose is very conservative. Our results suggest, however, that it is still effective. Regulators (APVMA) also limited the number of treatment baits available at a site at any one time to 30 baits per hectare. It is clear from our study that, at this density of baits, many quolls are simply not encountering the bait; arriving at the bait station after baits have been taken; this in a relatively low density quoll population, and despite multiple bait events at each site. Thus, to effectively bait a large proportion of the quolls at a site (particularly a high density site), a greater density of baits will be required.

In addition to the high visitation rate of individual quolls to bait stations, some individual quolls took baits on multiple occasions. Of the 70 “individual” quolls that visited bait stations throughout study period, ten individuals consumed a cane toad sausage on more than one occasion within a session. Why did these individuals manifestly fail to train? One possibility is the low dose rate, 0.06g of TBZ in each sausage was calculated to provide 300mg/kg to the smallest adult quoll at our site; a 200g female. Long-term trapping at the site (AWC, unpub. data) suggests that adult quolls in this population can reach more than 815g in weight. Thus, large animals could receive less than one quarter of the dose ingested by small animals. As a consequence, we could expect larger animals (typically males) to be harder to train with a fixed-dose bait. Another possibility is that these individuals were unhealthy for other reasons (e.g., males in the process of annual die-off) and so were willing to risk poisoning in order to acquire food, although such a mechanism would presumably cause changes in uptake rate across sessions, so seems unlikely.

Our results also hint strongly that individuals lose their acquired aversion over the 4-5 month window between our baiting sessions. There was no evidence that first pass rates of bait uptake declined over time across sessions. Whether this aversion would decline in the presence of ongoing stimulus (i.e., continuous baiting, or the presence of toads) is unknown, but long-term mark-recapture studies of CTA-trained quolls released into toad-infested landscapes suggest that aversion can be long-held in the presence of reinforcing stimulus (Cremona *et al.* 2017). Nonetheless, our finding should sound a note of caution with regard to deployment of CTA. Training prior to toad arrival will need to be delicately timed: too early, and trained animals may lose their aversion before toads arrive. This need for precision timing is complicated by inevitable uncertainty with regard to where the toad invasion front is, and when it will arrive at the site (with spread rate also being contingent on the unpredictable timing of the wet season in northern Australia). Thus, any baiting campaign will need to dedicate effort to predicting the date of toad arrival at the site.

Overall, however, our study is encouraging with regard to the use of toad sausages as a vehicle for large-scale CTA training of quolls. Our results suggest that quolls will consume cane toad sausages in the field and will, as a consequence, be less inclined to attack cane toads. This opens the possibility for broad scale application of CTA as a management technique for mitigating the impact of toads on quolls.

While many questions remain about optimal bait design, delivery, and timing, it is clear that CTA training using toad sausages is likely a viable tool for land managers seeking to protect quoll populations. Given that quoll populations in the Kimberley will likely be completely overrun by toads within the next five years, this is a tool that is urgently needed. More generally, however, our work joins a growing list of studies demonstrating that the impact of invasive species can be mitigated not only by controlling the invasive species, but also – or instead - by manipulating its mechanism of impact.

## Authors’ Contributions

All persons who meet authorship criteria are listed as authors, and all authors certify that they have participated sufficiently in the work to take public responsibility for the content, including participation in the concept, design, analysis, writing, or revision of the manuscript.

## Acknowledgements

We thank Australian Wildlife Conservancy and their supporters for access to northern quolls and their field laboratory. We also wish to acknowledge the support of the Territory Wildlife Park for assistance with the captive trials and with housing and husbandry of the quolls in captivity. Many thanks to Kristina Koenig, whose efforts assisted us immensely with data collation and individual identification of quolls. Additionally we wish to thank our volunteers who participated with field trials.

Funding to undertake this research was provided by Australian Wildlife Conservancy supporters, the Australian Research Council through LP150100722 and FT160100198 to B.L.P. and J. Webb). Holsworth Wildlife Research Endowment (to N.I.) and The Australian Wildlife Society-Wildlife Ecology Science Research Scholarship (to N.I.).

## Data accessibility

Data and scripts are available upon request.

## Compliance with Ethical Standards

The study was conducted within Mornington Wildlife Sanctuary in accordance with Wildlife Conservation Regulation 17 (Permit number: SFO10584). The area is jointly managed by traditional land-owners and the Australian Wildlife Conservancy. The research was approved by the University of Melbourne Animal Ethics Committee (Protocol: 1413369.2) and the University of Technology Sydney Animal Care and Ethics Committee (Protocol: 2012-432A) and Department of Parks and Wildlife Animal Ethics Committee (Protocol: DPaW AEC 2016_50 and Protocol 2013_37). Additionally this study was conducted in accordance with the approved outline submitted to the AVPMA by the investigating team (Permit to allow the possession and supply for research use of an unregistered Agvet chemical product. Permit number: PER92262).

## References

Avery, M.L. (1985) Application of Mimicry Theory to Bird Damage Control. The Journal of Wildlife Management, 49, 1116–1121.

Avery, M.L., Pavelka, A.A., Bergman, D.L., Decker, D.G., Kinttle, C.E. & Linz, G.M. (1995) Aversive conditioning to reduce raven predation on California least tern eggs. Journal of the Colonial Waterbird Society, 18, 131–245.

Bernstein, I.L. (1999) Taste aversion learning: a contemporary perspective. Nutrition, 15, 229–234.

Bogliani, G. & Fiorella, B. (1998) Conditioned Aversion as a Tool to Protect Eggs from Avian Predators in Heron Colonies. Colonial Waterbirds, 21, 69–72.

Bolger, D.T., Vance, B.T., Morrison, T.A. & Farid, H. (2011) Wild-ID. Dartmouth College, Hanover, NH.

Burnett, S. (1997) Colonizing Cane Toads cause population declines in native predators: reliable anecdotal information and management implications. Pacific Conservation Biology, 3, 65–72.

Claridge, A.W. & Mills, D.J. (2007) Aerial baiting for wild dogs has no observable impact on spotted-tailed quolls (*Dasyurus maculatus*) in a rainshadow woodland. Wildlife Research, 34, 116–124.

Cohn, J. & MacPhail, R.C. (1996) Ethological and Experimental Approaches to Behavior Analysis: Implications for Ecotoxicology. Environmental Health Perspectives, 104, 299–305.

Conover, M.R. (1989) Potential compounds for establishing conditioned food aversions in Racoons. Wildlife Society Bulletin, 17.

Conover, M.R. (1995) Behavioral Principles Governing Conditioned Food Aversions Based on Deception. USDA National Wildlife Research Center Symposia National Wildlife Research Center Repellents Conference pp. 29–41. DigitalCommons@University of Nebraska - Lincoln, University of Nebraska - Lincoln.

Covacevich, J. & Archer, M. (1975) The distribution of the cane toad, *Bufo marinus*. Australia and its effects on indigenous vertebrates. Memoirs of the Queensland Museum, 17, 305–310.

Cowled, B.D., Gifford, E., Smith, M., Staples, L. & Lapidge, S.J. (2006) Efficacy of manufactured PIGOUT® baits for localised control of feral pigs in the semi-arid Queensland rangelands. Wildlife Research, 33, 427–437.

Cox, R., Baker, S.E., Macdonald, D.W. & Berdoy, M. (2004) Protecting egg prey from Carrion Crows: the potential of aversive conditioning. Applied Animal Behaviour Science, 87, 325–342.

Cremona, T., Spencer, P., Shine, R. & Webb, J.K. (2017) Avoiding the last supper: parentage analysis indicates multi-generational survival of re-introduced ‘toad-smart’lineage. Conservation Genetics, 1–6.

Diete, R.L., Meek, P.D., Dixon, K.M., Dickman, C.R. & Leung, L.K.-P. (2016) Best bait for your buck: bait preference for camera trapping north Australian mammals. Australian Journal of Zoology, 63, 376–382.

Dilov, P., Chaleva, E., Dzhurov, A., Stoianov, P. & Iotsev, M. (1981) [Toxicological evaluation of “Farmakhim” thiabendazole in mammals and birds]. Veterinarnomeditsinski nauki, 19, 92–100.

Doherty, T.S., Dickman, C.R., Glen, A.S., Newsome, T.M., Nimmo, D.G., Ritchie, E.G., Vanak, A.T. & Wirsing, A.J. (2017) The global impacts of domestic dogs on threatened vertebrates. Biological Conservation, 210, Part A, 56–59.

Dundas, S., Adams, P. & Fleming, P. (2014) First in, first served: uptake of 1080 poison fox baits in south-west Western Australia. Wildlife Research, 41, 117–126.

Ellins, S.R. & Catalano, S.M. (1980) Field application of the conditioned taste aversion paradigm to the control of coyote predation on sheep and turkeys. Behavioral and Neural Biology, 29, 532–536.

Fairbridge, D., Anderson, R., Wilkes, T. & Pell, G. (2003) Bait uptake by free living brush-tailed phascogales Phascogale tapoatafa and other non-target mammals during simulated buried fox baiting. Australian Mammology, 25, 31–40.

Gill, E.L., Whiterow, A. & Cowan, D.P. (2000) A comparitive assessment of potential conditioned taste aversion agents for vertebrate management. Applied Animal Behaviour Science, 67, 229–240.

Glen, A.S. & Dickman, C., R (2003) Monitoring bait removal in vertebrate pest control: a comparison using track identification and remote photography. Wildlife Research, 30, 29–33.

Glendinning, J.I. (2007) How Do Predators Cope With Chemically Defended Foods?. The Biological Bulletin, 213, 252–266.

Gustavson, C.R. (1982) An evaluation of taste aversion control of wolf (*Canis lupus*) predation in Northern Minnesota. Applied Animal Ethology, 9, 63–71.

Gustavson, C.R. & Basche, L.A. (1983) Landrin-and thiabendazole-based conditioned taste aversions in domestic ducks. Applied Animal Ethology, 9, 379–380.

Gustavson, C.R., Garcia, J., Hankins, W.G. & Rusiniak, K.W. (1974) Coyote Predation Control by Aversive Conditioning. Science, 184, 581–583.

Gustavson, C.R., Gustavson, J.C. & Holzer, G.A. (1983) Thiabendazole-based taste aversions in dingoes (*Canis familiaris dingo*) and New Guinea wild dogs (*Canis familiaris hallstromi*). Applied Animal Ethology, 10, 385–388.

Gustavson, C.R., Kelly, D.R., Sweeney, M. & Garcia, J. (1976) Prey-Lithium Aversions. I: Coyotes and Wolves. Behavioral Biology, 17, 61–72.

Gustavson, C.R. & Nicolaus, L.K. (1987) Taste aversion conditioning in wolves, coyotes, and other canids: retrospect and prospect. Man and wolf, 169–200.

Hayes, R.A., Crossland, M.R., Hagman, M., Capon, R.J. & Shine, R. (2009) Ontogenetic Variation in the Chemical Defenses of Cane Toads (*Bufo marinus*): Toxin Profiles and Effects on Predators. Journal of Chemical Ecology, 35, 391–399.

Hohnen, R., Ashby, J., Tuft, K. & McGregor, H. (2013) Individual identification of northern quolls (Dasyurus hallucatus) using remote cameras. Australian Mammalogy, 35, 131–135.

Jolley, W.J., Campbell, K.J., Holmes, N.D., Garcelon, D.K., Hanson, C.C., Will, D., Keitt, B.S., Smith, G. & Little, A.E. (2012) Reducing the impacts of leg hold trapping on critically endangered foxes by modified traps and conditioned trap aversion on San Nicolas Island, California, USA. Conservation Evidence, 9, 43–49.

Jolly, C.J., Shine, R. & Greenlees, M.J. (2015) The impact of invasive cane toads on native wildlife in southern Australia. Ecology and Evolution, 5, 3879–3894.

Jolly et al. unpublished data.

Kelly, E. & Phillips, B. (2017) Get smart: native mammal develops toad-smart behavior in response to a toxic invader. Behavioral Ecology, 28, 854–858.

Legge, S., Murphy, B.P., McGregor, H., Woinarski, J.C.Z., Augusteyn, J., Ballard, G., Baseler, M., Buckmaster, T., Dickman, C.R., Doherty, T., Edwards, G., Eyre, T., Fancourt, B.A., Ferguson, D., Forsyth, D.M., Geary, W.L., Gentle, M., Gillespie, G., Greenwood, L., Hohnen, R., Hume, S., Johnson, C.N., Maxwell, M., McDonald, P.J., Morris, K., Moseby, K., Newsome, T., Nimmo, D., Paltridge, R., Ramsey, D., Read, J., Rendall, A., Rich, M., Ritchie, E., Rowland, J., Short, J., Stokeld, D., Sutherland, D.R., Wayne, A.F., Woodford, L. & Zewe, F. (2017) Enumerating a continental-scale threat: How many feral cats are in Australia? Biological Conservation, 206, 293–303.

Letnic, M., Webb, J.K. & Shine, R. (2008) Invasive cane toads (*Bufo marinus*) cause mass mortality of freshwater crocodiles (*Crocodylus johnstoni*) in tropical Australia. Biological Conservation, 141, 1773–1782.

Llewelyn, J., Phillips, B.L., Brown, G.P., Schwarzkopf, L., Alford, R.A. & Shine, R. (2011) Adaptation or preadaptation: why are keelback snakes (Tropidonophis mairii) less vulnerable to invasive cane toads (Bufo marinus) than are other Australian snakes? Evolutionary Ecology, 25, 13–24.

Mappes, J., Marples, N. & Endler, J. (2005) The complex business of survival by aposematism. Trends in Ecology and Evolution, 20, 598–603.

Massei, G. & Cowan, D.P. (2002) Strength and persistence of conditioned taste aversion in rats: evaluation of 11 potential compounds. Applied Animal Behaviour Science, 75, 249–260.

Massei, G., Lyon, A. & Cowan, D.P. (2003) Levamisole can induce conditioned taste aversion in foxes. Wildlife Research, 30, 633–637.

Nachman, M. & Ashe, J. (1973) Learned Taste Aversions in Rats as a Function of Dosage, Concentration, and Route of Administration of LiCl. Physiology and Behavior, 10, 73–78.

Nicolaus, L.K. (1987) Conditioned Aversions in a guild of egg predators: Implications for aposematism and prey defense mimicry. American Midland Naturalist, 117, 405–419.

Nicolaus, L.K., Herrera, J., Nicolaus, J.C. & Dimmick, C.R. (1989) Carbachol as a conditioned taste aversion agent to control avian depredation. Agriculture, Ecosystems and Environment, 26, 13–21.

Nicolaus, L.K. & Nellis, D.W. (1987) The First Evaluation of the Use of Conditioned Taste Aversion to Control Predation by Mongooses upon Eggs. Applied Animal Behaviour Science, 17, 329–346.

O’Donnell, S., Webb, J.K. & Shine, R. (2010) Conditioned taste aversion enhances the survival of an endangered predator imperilled by a toxic invader. Journal of Applied Ecology, 47, 558–565.

Oakwood, M. & Foster, P. (2008) Monitoring extinction of the northern quoll. Australian Academy of Science Newsletter, 71,

Page, R.A. & Ryan, M.R. (2005) Flexibility in assessment of prey cues: frog-eating bats and frog calls. Proceedings of the Royal Society, 272, 841–847.

Phillips, B.L. & Shine, R. (2005) The morphology, and hence impact, of an invasive species (the cane toad, Bufo marinus): changes with time since colonisation. Animal Conservation, 8, 407–413.

Provenza, F.D., Burritt, E.A., Clausen, T.P., Bryant, J.P., Reichardt, P.B. & Distel, R.A. (1990) Conditioned flavor aversion: a mechanism for goats to avoid condensed tannins in blackbrush. American Naturalist, 810–828.

Reaser, J.K., Meyerson, L.A., Cronk, Q., De Poorter, M., Eldrege, L.G., Green, E., Kairo, M., Latasi, P., Mack, R.N. & Mauremootoo, J. (2007) Ecological and socioeconomic impacts of invasive alien species in island ecosystems. Environmental Conservation, 34, 98–111.

Riley, A.L. & Freeman, K.B. (2004) Conditioned taste aversion: a database. Pharmacology Biochemistry and Behavior, 77, 655–656.

Risbey, D.A., Calver, M.C., Short, J., Bradley, J.S. & Wright, I.W. (2000) The impact of cats and foxes on the small vertebrate fauna of Heirisson Prong, Western Australia. II. A field experiment. Wildlife Research, 27, 223–235.

Robinson, H.J., Stoerk, H.C. & Graessle, O.E. (1965) Studies on the toxicologic and pharmacologic properties of thiabendazole. Toxicology and applied pharmacology, 7, 53–63.

Schneider, K. & Pinnow, M. (1994) Olfactory and Gustatory Stimuli in Food-Aversion Learning of Rats. The Journal of General Psychology, 121, 169–183.

Semel, B. & Nicolaus, L.K. (1992) Estrogen-Based Aversion to Eggs Among Free-Ranging Raccoons. Ecological Applications, 2, 439–449.

Sevelinges, Y., Mouly, A.M., Levy, F. & Ferreira, G. (2009) Long-Term Effects of Infant Learning on Adult Conditioned Odor Aversion Are Determined by the Last Preweaning Experience. Developmental Psychobiology, 299–398.

Shine, R. (2010) The ecological impact of invasive cane toads (*Bufo marinus*) in Australia. The Quarterly Review Of Biology, 85, 253–291.

Short, J. & Smith, A. (1994) Mammal decline and recovery in Australia. Journal of Mammalogy, 75, 288–297.

Simberloff, D., Martin, J.-L., Genovesi, P., Maris, V., Wardle, D.A., Aronson, J., Courchamp, F., Galil, B., García-Berthou, E., Pascal, M., Pyšek, P., Sousa, R., Tabacchi, E. & Vilà, M. (2013) Impacts of biological invasions: what's what and the way forward. Trends in Ecology & Evolution, 28, 58–66.

Sinclair, R.G. & Bird, P.L. (1984) The reaction of Sminthopsis crassicaudata meat baits containing 1080: Implications for assessing risk to non-target species. Australian Wildlife Research, 11, 501–507.

Smith, J. & Phillips, B.L. (2006) Toxic tucker: the potential impact of Cane Toads on Australian reptiles. Pacific Conservation Biology, 12, 40–49.

Smith, M.E., Linnell, J.D.C., Odden, J. & Swenson, J.E. (2000) Review of Methods to Reduce Livestock Depredation II. Aversive conditioning, deterrents and repellents. Acta Agriculturae Scandinavica, Section A — Animal Science, 50, 304–315.

Ternent, M.A. & Garshelis, D.L. (1999) Taste-aversion conditioning to reduce nuisance activity by black bears in a Minnesota military reservation. Wildlife Society Bulletin, 27, 720–728.

Thornton, A. & Clutton-Brock, T. (2011) Social learning and the development of individual and group behaviour in mammal societies. Philosophical Transactions of the Royal Society B: Biological Sciences, 366, 978–987.

Thornton, A. & Raihani, N.J. (2010) Identifying teaching in wild animals. Learning & Behavior, 38, 297–309.

Tocco, D.J., Rosenblum, C., Martin, C.M. & Robinson, H.J. (1966) Absorption, metabolism, and excretion of thiabendazole in man and laboratory animals. Toxicology and applied pharmacology, 9, 31–39.

Urban, M.C., Phillips, B.L., Skelly, D.K. & Shine, R. (2007) The cane toad’s (*Chaunus [Bufo] marinus*) increasing ability to invade Australia is revealed by a dynamically updated range model. Proceedings of the Royal Society Biological Sciences Series B, 274, 1413–1419.

Webb, J.K., Brown, G.P., Child, T., Greenlees, M.J., Phillips, B.L. & Shine, R. (2008) A native dasyurid predator (common planigale, *Planigale maculata*) rapidly learns to avoid a toxic invader. Austral Ecology, 33, 821–829.

Webb, J.K., Pearson, D. & Shine, R. (2011) A small dasyurid predator (*Sminthopsis virginiae*) rapidly learns to avoid a toxic invader. Wildlife Research, 38, 726–731.

Webb, J.K., Shine, R. & Christian, K.A. (2005) Does intraspecific niche partitioning in a native predator influence its response to an invasion by a toxic prey species? Austral Ecology, 30, 201–209.

Woinarski, J., Burbidge, A. & Harrison, P. (2014) The action plan for Australian mammals 2012.

Ziegler, J.M., Gustavson, C.R., Holzer, G.A. & Gruber, D. (1983) Anthelmintic-based taste aversions in wolves (*Canis lupus*). Applied Animal Ethology, 9, 373–377.

Ziembicki, M.R., Woinarski, J.C.Z., Webb, J.K., Vanderduys, E., Tuft, K., Smith, J., Ritchie, E.G., Reardon, T.B., Radford, I.J., Preece, N., Perry, J., Murphy, B. P., McGregor, H., Legge, S., Leahy, L., Lawes, M.J., Kanowski, J., Johnson, C.N., James, A., Griffiths, A.D., Gillespie, G., Frank, A.S.K., Fisher, A. & Burbidge, A.A. (2015) Stemming the tide: progress towards resolving the causes of decline and implementing management responses for the disappearing mammal fauna of northern Australia. Therya, 6, 169–225.

